# Exploring Organ-Specific Extracellular Vesicles in Metabolic Improvements Following Bariatric Surgery in Adolescents with Obesity

**DOI:** 10.1101/2024.11.07.622509

**Authors:** Ahlee Kim, Tsuyoshi Okura, Kwangmin Choi, Rupinder Gill, Vishnupriya J Borra, Kazutoshi Murakami, Andrew Poulos, Xiang Zhang, Todd Jenkins, Amy Sanghavi Shah, Michael Helmrath, Takahisa Nakamura

## Abstract

**Aims/hypothesis:** Vertical sleeve gastrectomy (VSG) leads to significant metabolic improvements, though the underlying molecular mechanisms are not yet fully understood. Emerging evidence suggests that small extracellular vesicles (sEVs) contribute to metabolic improvements post-VSG; however, it is still unclear which organ-specific sEV correlate with various metabolic parameters and how they exert these effects. The study aimed to establish the role of organ-specific sEVs in the metabolic improvements associated with VSG.

**Methods:** Demographic, anthropometric, and blood samples were collected pre-VSG and 3- and 6-month post-VSG in adolescents with obesity. Blood samples were utilized to measure metabolic parameters and to isolate sEVs. sEV RNAs were analyzed via small RNA sequencing then bioinformatics analyses.

**Results:** A significant reduction in mRNA cargo from liver-specific genes was observed post-VSG, whereas adipose tissue- or skeletal muscle-specific genes showed no such reduction. Liver-derived RNA correlated with BMI, leptin, and resistin, while adipose-derived RNA correlated with leptin. Analysis of delta values (post-minus pre-surgery) revealed that adipose-derived RNA cargo correlated with markers of liver damage and HOMA-IR, whereas liver-derived RNA cargo correlated with BCAAs.

**Conclusions:** VSG modulates the EV system in the liver and adipose tissue. Liver-derived sEVs appear to regulate adipose metabolism, while adipose-derived sEVs are associated with liver function, suggesting a dynamic crosstalk between these tissues through sEVs that shapes systemic metabolic outcomes.

**Research in Context:** *What is already known about this subject?:*

- Vertical sleeve gastrectomy (VSG) is the most performed bariatric surgery, leading to significant metabolic improvements.
- Small extracellular vesicles (sEVs) and their RNA cargo play a crucial role in metabolic regulation.
- Levels of branched-chain amino acids (BCAAs), which are known metabolic regulators, decrease following bariatric surgery.

*What is the key question?:*

- How do sEVs, BCAA and other metabolic molecules interact to drive the significant metabolic improvements observed after VSG?

*What are the new findings?:*

- The levels of liver-specific RNA cargo in sEVs decrease after VSG and are correlated with BMI and leptin levels at 6 months post-surgery, suggesting a potential role for liver-specific RNA cargo in the metabolic benefits associated with VSG.
- Changes in BCAA levels from baseline to 6 months post-surgery correlate with changes in liver-specific RNA cargo in sEVs, indicating liver-specific RNA cargo in sEVs may influence metabolic parameters through BCAA modulation.
- Changes in the level of adipose-specific RNA cargo in sEVs from baseline to 6 month post-surgery correlate with changes HOMA-IR and ALT levels.

*How might this impact on clinical practice in the foreseeable future?:* The study identifies liver- and adipose-derived sEV RNA cargo as potential targets for non-invasive therapies that could replicate the metabolic benefits of VSG. While further investigation is needed, these findings suggest a tentative alternative for adolescents facing obesity who may be ineligible or reluctant to undergo bariatric surgery.

## Introduction

Obesity poses significant health risks, including insulin resistance, type 2 diabetes (T2D), hypertension, dyslipidemia, cardiovascular disease, and metabolic-dysfunction associated steatotic liver disease (MASLD) and metabolic-dysfunction associated steatotic hepatitis (MASH) [1]. Bariatric surgery is a safe and effective intervention to promote dramatic weight loss and metabolic improvements across age groups, including adults, adolescents, and children [1–3]. Specifically, vertical sleeve gastrectomy (VSG) is the most commonly performed bariatric surgery in adolescents, demonstrating both safety and effectiveness, with an average 26% weight loss reported 1-year post-VSG [4]. As a result, the current clinical guideline for pediatric obesity management now includes bariatric surgery as part of the standard of care [1, 2].

Without a doubt, one of the primary mechanisms behind metabolic improvements associated with VSG is significant weight loss [2, 3, 5–7]. However, accumulating evidence suggests that these metabolic benefits are not entirely dependent on weight loss alone. Remarkably, metabolic changes appear within just a few days of surgery, well before substantial weight loss occurs, and persist even when some weight is regained [6, 8–11]. These findings have sparked interest in identifying the molecular mechanisms underlying the weight-loss-independent benefits of bariatric surgery.

While data for levels of cytokines, including adipokines, following bariatric surgery, remains inconsistent [12], some adipokines, such as leptin and resistin, are thought to contribute to metabolic improvements associated with bariatric surgery [13]. For instance, levels of both leptin and resistin decrease after surgery [14]. The reduction in leptin correlates strongly with changes in body mass index (BMI), suggesting that the decrease in fat mass and improvement in leptin resistance are interconnected [15]. Resistin levels, similarly, are closely associated with weight and BMI, with a downward trend observed post-surgery, highlighting its role in obesity-related metabolic complications [16]. In addition to adipokines, branched-chain amino acids (BCAAs), a subgroup of amino acids with an aliphatic side-chain that includes leucine, isoleucine, and valine [17], have emerged as key regulators of metabolic health and are crucial for human metabolism [18–20], particularly in the context of obesity, insulin resistance, and T2D [18]. Elevated BCAA levels are consistently linked to metabolic dysfunction, including impaired glucose metabolism. Notably, recent studies indicate that bariatric surgery significantly lowers circulating BCAA levels, contributing to the observed metabolic improvements [21].

Small extracellular vesicles (sEVs) carry various molecules, including genetic materials, between tissues, acting as key mediators of inter-organ communication [22–24]. Small EVs are associated with a range of metabolic conditions, including but not limited to insulin resistance, T2D, MASLD/MASH, and cardiovascular disease [25–32]. Recent studies suggest that the metabolic benefits of bariatric surgery involve alterations in the circulating sEV profile [33]. Studies have shown that fatty acid binding protein 4 carried by sEVs increases 1 month after bariatric surgery and returns to baseline by 6 months post-surgery [34]. Another investigation found a reduction in hepatocyte-specific sEVs following bariatric surgery in adults [35]. Similarly, research has reported a decrease in sEVs containing hepatocyte-specific surface markers in adults with obesity after surgery [36].

Here we report changes in the sEV and metabolic parameters in adolescents with severe obesity undergoing VSG, and how these changes correlate with metabolic outcomes before and after VSG. In particular, we focused on the tissue-specific genes in circulating sEVs to understand how different organs are affected by VSG. Furthermore, we examined how tissue-specific sEV cargo interacts with BCAA in driving robust metabolic improvement following VSG.

## Methods

### Study Population

Adolescents with severe obesity undergoing VSG were recruited and enrolled as part of a single-center, multidisciplinary study at the Pediatric Diabetes and Obesity Center at Cincinnati Children’s Hospital Medical Center (CCHMC) from 2016 to 2019. The decision to proceed with VSG was made in accordance with the American Society for Metabolic and Bariatric Surgery (ASMBS) pediatric guidelines [2], involving the subjects, their families, and the multidisciplinary clinical team. All surgery was performed by a single pediatric bariatric surgeon at CCHMC. Participants were assessed and provided blood samples at prior to VSG, and 3- and 6-months post-VSG. Written informed consent was obtained from participants aged 18 years or older, or from a legal guardian for those under 18 years old, with assent obtained from the minors. The study was reviewed and approved by the Institutional Review Board (IRB) at CCHMC.

### Anthropometric Data

Height was measured in centimeters (cm) using a wall-mounted stadiometer, and weight was recorded in kilograms (kg) using an electronic scale. BMI was calculated as weight in kilograms divided by height in meters squared (kg/m^2^).

### Isolation of sEVs and Extraction of Small RNA Cargo

Small EVs were isolated from serum samples using the MagCapture™ Exosome Isolation Kit (Wako Pure Chemical Corporation, Osaka, Japan), which utilizes Tim4 protein immobilized on magnetic beads to bind phosphatidylserine exposed on the surface of sEVs. A volume of 500 µL to 1 mL of serum was used for each isolation. The serum was incubated with the Tim4-coated magnetic beads on a rotator at room temperature for 30 minutes, allowing efficient capture of EVs. Following incubation, the magnetic beads were separated using a magnetic stand, and unbound components were removed through several washes with phosphate-buffered saline. The EVs were then eluted from the beads with the provided elution buffer and immediately used for RNA extraction. For RNA isolation, the Qiagen miRNeasy Serum/Plasma Kit (Qiagen, Hilden, Germany) was employed according to the manufacturer’s protocol. Small RNAs, including miRNAs, were extracted from sEVs isolated from the same amount of serum across all samples to ensure consistency. The quality and concentration of the isolated RNA were assessed before proceeding to downstream molecular analyses, such as small RNA sequencing.

### Small EV RNA cargo sequencing

Small RNA-seq of RNA cargo was performed by the Genomics, Epigenomics, and Sequencing Core at the University of Cincinnati [37, 38]. For library preparation, the NEBNext Small RNA Sample Library Preparation Kit (NEB, Ipswich, MA) was used with a modified approach for precise small RNA library size selection, allowing the kit to process low-input RNA (as low as ng levels) with improved library recovery. Specifically, 0.4–4 ng of sEVs RNA, quantified by Bioanalyzer QC, was used as input. Following 15 cycles of PCR, libraries with unique indices were pooled at equal volumes (10 µl per sample), cleaned using a column-based approach, and mixed with a custom-designed DNA ladder containing 135 bp and 319 bp PCR amplicons. This size range corresponds to small RNA libraries with 16–200 nt inserts, encompassing all small RNA species, including miRNAs. After high-resolution agarose gel electrophoresis and gel excision, the library pool—ranging from 135 to 319 bp, including the custom DNA ladder—was purified and quantified using the NEBNext Library Quant Kit on a QuantStudio 5 Real-Time PCR System (ThermoFisher, Waltham, MA). An initial round of sequencing was performed on a NextSeq 550 sequencer (Illumina, San Diego, CA) to generate several million reads, enabling the relative quantification of each library. The volume of each library was then adjusted, followed by sequencing with single-end 1×85 bp reads to generate ∼17 million reads per sample for final data analysis.

### Sequencing Data Preparation and Downstream Analyses

Sequencing reads were quality-checked using FASTQC (v. 0.11.2; https://www.bioinformatics.babraham.ac.uk/projects/fastqc/). The 3’ adapter sequence (“AGATCGGAAGAGCACACGTCTGAACTCCAGTCAC”) was clipped, and reads were trimmed using the FASTX toolkit (v. 0.0.14; http://hannonlab.cshl.edu/fastx_toolkit/). Clipped reads were quantified with the excerpt tool (v. 4.6.2) on the Genboree Workbench (http://www.genboree.org/). First, reads were aligned using Bowtie2 to UniVec, a library of common contaminant sequences maintained by the NCBI, with valid alignments archived and removed from further analysis. Next, reads were aligned to human ribosomal RNA (rRNA) precursor sequences (including the full 45S, 5S, and mitochondrial rRNAs) using Bowtie2. Reads that passed these filters were aligned sequentially to several databases: Human genome (hg19), miRNA (miRBase; http://www.mirbase.org/), tRNA (gtRNAdb; http://gtrnadb.ucsc.edu/), piRNA (piRNABank; http://pirnabank.ibab.ac.in/), long-RNA (GENCODE; https://www.gencodegenes.org/), and circRNA (circBase; http://www.circbase.org/). Aligned read counts were normalized using the TMM (Trimmed Mean of M-values) method implemented in the edgeRpackage (v. 3.10; https://bioconductor.org/packages/release/bioc/html/edgeR.html). Additionally, downsampling was applied to handle differences in library sizes across samples and batches. Heatmaps and PCA plots were generated using the pheatmap (https://cran.r-project.org/web/packages/pheatmap/index.html) and ggplot2 (https://ggplot2.tidyverse.org/) packages, respectively. Tissue-specific and tissue-enriched genes were collected from the TissueEnrich database’s Human Protein Atlas (https://tissueenrich.gdcb.iastate.edu/). For statistical testing, the non-parametric Kruskal-Wallis test by ranks was conducted with R (v4.4.0) to evaluate whether significant differences exist in RNA and clinical marker levels among paired samples at 0, 3, and 6 months. Correlations between RNA and clinical markers were assessed using R (v4.4.0), and scatter plots were generated with the ggplot2 package (v3.5.2).

### Clinical Labs

Blood samples were collected via venipuncture after participants completed an overnight fast (>8 hours). Fasting glucose level was measured using a Hitachi model 704 glucose analyzer (Roche Hitachi, Indianapolis, IN) and fasting insulin levels were assessed by radioimmunoassay with anti-insulin serum raised in guinea pigs, ^125^I-labeled insulin (Linco, St Louis, MO) and a double antibody method to separate bound from free tracer. The Homeostatic Model Assessment of Insulin Resistance (HOMA-IR) was calculated using as: HOMA-IR = (Fasting glucose (mg/dL) × Fasting insulin (µU/mL)) / 405. Alanine aminotransferase (ALT) was measured using a Siemens Atellica clinical analyzer (Siemens Healthineers, Erlangen, Germany). Leptin and resistin levels were measured using the Luminex assay platform (Luminex Corporation, Austin, TX) at CCHMC, following the manufacturer’s protocol.

### BCAA measurement

BCAAs were quantified using a Thermo Scientific Vanquish UHPLC system coupled to a Q Exactive Plus Hybrid Quadrupole-Orbitrap Mass Spectrometer (Thermo Fisher Scientific). Chromatographic separation was performed on an ACE C18-PFP column (100 mm × 2.1 mm) at 30°C, with a flow rate of 0.2 mL/min. The mobile phase consisted of water with 0.1% formic acid (FA) (solvent A) and acetonitrile with 0.1% FA (solvent B). A gradient reduced solvent A from 99% to 10% over 8 minutes, followed by re-equilibration. For serum sample preparation, 25 µL of thawed serum was mixed with 100 µL of internal standard solution (5 µg/mL D8-valine and D10-leucine in acetonitrile), vortexed, and centrifuged at 21,000g and 4°C for 15 minutes. The supernatant was dried under nitrogen at 35°C, reconstituted in 300 µL LC/MS-grade water with 0.1% FA, and 5 µL was injected for analysis. Calibration standards (0.5–100 µg/mL) and quality control samples (1, 20, 40, 80 µg/mL) were prepared in 0.1% FA. The mass spectrometer operated in positive electrospray ionization mode, with MS2 scans at 70,000 resolutions. Precursor ions of leucine, isoleucine, and valine were quantified using targeted transitions: m/z 118.0865 → 55.0552 for valine and m/z 132.1019 → 86.0970 for leucine and isoleucine. Data were processed using Xcalibur 4.1, with peak areas extracted within a 10 ppm mass window. The assay was linear across a range of 0.5–100 µg/mL.

## Results

### Baseline Characteristics of the Cohort

The study included 58 adolescents with obesity, with a mean age of 16.7 ± 1.7 years and a mean BMI of 50.6 ± 9.6 kg/m², categorizing them as class 3 obesity prior to VSG. Of these participants, 69% were female. **Table 1** provides detailed baseline characteristics, including height, weight, and key metabolic parameters such as glucose and insulin levels, indicating the presence of insulin resistance. Liver function was assessed through ALT levels, and BCAA levels were also measured to evaluate metabolic health. Of note, BCAA levels were measured in 45 samples due to sample limitations.

**Table 1.** Baseline Characteristics of Study Subjects. A table presenting the baseline characteristics of adolescents with obesity who participated in the study prior to undergoing VSG.

| Baseline Parameter | Mean |
| --- | --- |
| n | 58 |
| Male/Female | 18/40 |
| Age (years) | 16.7 +/- 1.7 |
| Height (cm) | 169.3 +/- 10.2 |
| Body weight (kg) | 145.3 +/-29.6 |
| BMI (kg/m2) | 50.6 +/- 9.6 |
| Glucose (mmol/L) | 5.18 +/- 1.53 |
| Insulin (pmol/L) | 69.2 +/- 34.3 |
| HOMA-IR | 2.65 +/- 0.38 |
| ALT (IU/L) | 47.2 +/- 45.1 |

### Anthropometric and Metabolic Parameters Post-VSG

Anthropometric and metabolic parameters were evaluated at multiple time points following VSG to assess the surgery’s impact on key metabolic markers. As previously reported, VSG led to a statistically significant reduction in BMI at both 3- and 6-months post-surgery [4, 8] (Fig. 1a). This result demonstrates that the weight loss trajectory established shortly after surgery is maintained, reflecting effective management of obesity through VSG. In addition to BMI reductions, significant changes were observed in metabolic hormones. Serum leptin levels, which are primarily secreted by adipose tissue and play a role in energy homeostasis [39], were significantly reduced at both 3 and 6 months post-VSG (Fig. 1b). This reduction correlates with decreased fat mass, as leptin levels are closely linked to adiposity. Serum resistin levels, a hormone implicated in inflammation and insulin resistance [40], showed a downward trend following VSG, although the decrease did not reach statistical significance (Fig. 1c). This variability may reflect individual differences in inflammatory responses or may require a longer follow-up period to observe clear effects.

**Figure 1.**
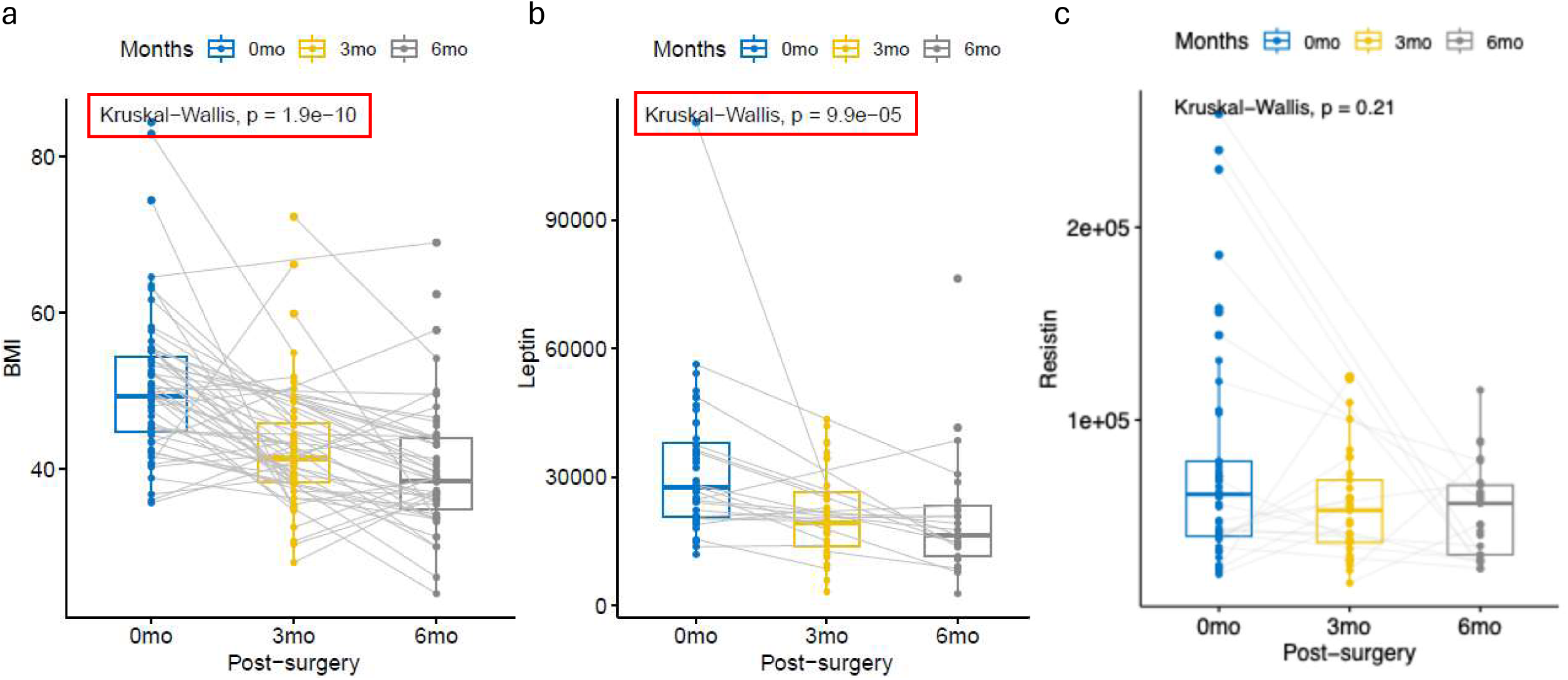
Changes in BMI, Leptin, and Resistin post-VSG. (a-c) Body Mass Index (BMI) (a), circulating Leptin (b), and Resistin (c) levels in obese adolescents measured pre-surgery (n=58), 3 months (n=44), and 6 months (n=40) post-VSG.

### Serum sEV Analysis and RNA Cargo Profiling

To explore whether sEVs are associated with the metabolic improvements observed post-VSG, we isolated sEVs from the serum of patients at each time point (Fig. 2a, 2b). Although serum sEV levels showed a downward trend post-VSG, the reduction was not statistically significant within the 6-month follow-up period (Fig. 2c). RNA cargo was further analyzed by isolating it from the collected sEVs, followed by RNA cargo-seq profiling. Among the RNA cargo, mRNA cargo was examined specifically to profile its tissue of origin. The RNA cargo profile exhibited a gradual shift from pre-surgery to 3- and 6-months post-VSG (Fig. 2d). Using the Human Protein Atlas (HPA) database, we classified these mRNAs based on tissue specificity (Supplemental Table 1). Notably, the abundance of liver-specific mRNA cargo (Liver-RNA cargo) was significantly reduced post-VSG (Fig. 2e), suggesting that the liver’s RNA cargo profile is highly sensitive to the metabolic changes induced by the surgery. In contrast, this pattern was not observed for RNA cargo derived from other key metabolic tissues, such as adipose tissue (Fig. 2f) and skeletal muscle (Fig. 2g and Supplemental Fig. 1). These findings indicate that obesity quantitatively modulates the RNA cargo profile of circulating sEVs, with the liver emerging as one of the most responsive organs to obesity-related metabolic changes. The data highlight the potential role of liver-derived sEVs in systemic metabolic regulation and underscore the liver’s unique vulnerability to obesity.

**Figure 2.**
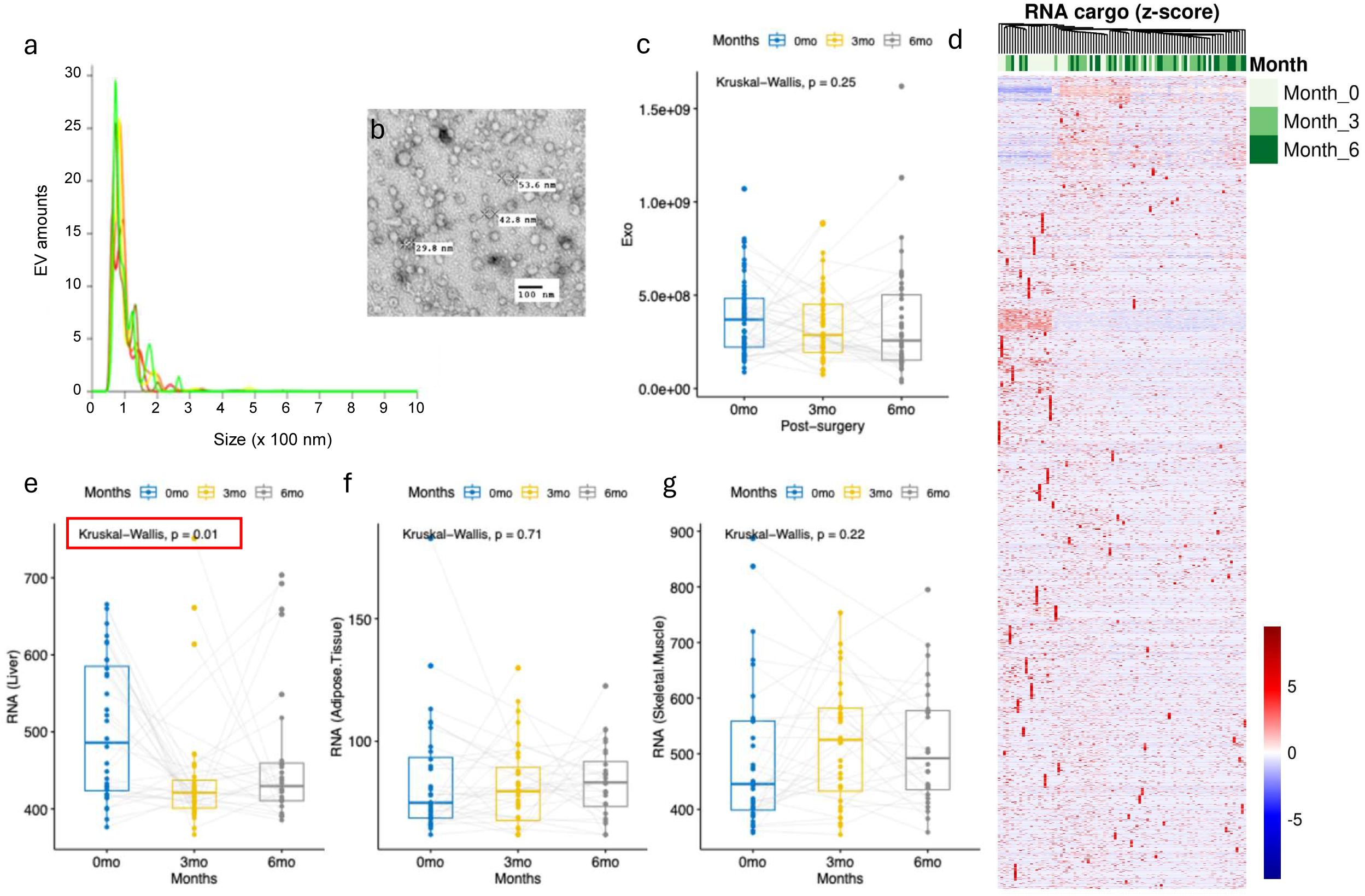
Effect of VSG on sEV RNA cargo profile. (a) Amount and size distribution of small extracellular vesicles (sEVs). (b) Transmission electron microscopy (TEM) image of circulating sEVs. (c) Concentration of small EVs in obese adolescents measured pre-surgery (n=58), 3-months (n=44), and 6-months (n=40) post-VSG. (d) Heat map depicting the mRNA cargo profile pre- and post-VSG. (e-g) RNA counts of tissue-specific genes within circulating sEVs for liver (e), adipose tissue (f), and skeletal muscle (g). Small RNA-seq read counts were generated using the Genboree pipeline. EdgeR was employed for read count normalization and to identify differentially expressed sEV-mRNA cargo. Tissue-enriched gene normalized counts were summed by tissue type, and combined abundances were plotted for pre-surgery, 3-month, and 6-month post-surgery time points.

### Post-VSG sEV-RNA Modulation and Metabolic Links

Given that the Liver-RNA cargo in circulating sEVs was significantly decreased post-VSG (Fig. 2c) and displayed trends similar to metabolic improvements, such as reductions in BMI (Fig. 1a), we next examined the correlation between Liver-RNA cargo and various metabolic parameters over time. A similar correlation analysis was also performed for other metabolic tissue-specific sEV-RNA levels, focusing on RNA cargo from adipose tissue-specific genes (Adipo-RNA cargo). At the pre-surgery time point, neither the Liver-RNA cargo or the Adipo-RNA cargo showed correlations with metabolic parameters such as BMI, leptin, or resistin levels (Supplemental Fig. 2a-2c). However, six months post-VSG, Liver-RNA cargo became positively correlated with both BMI and leptin (Fig. 3a, 3b), and, interestingly, it was negatively correlated with resistin (Fig. 3c), all with statistical significance. Additionally, Adipo-RNA cargo showed no pre-surgery correlations with BMI, leptin, or resistin (Supplemental Fig. 2d-2f), but it became negatively correlated with leptin 6 months post-VSG, reaching statistical significance (Fig. 2e), but not with BMI or resistin (Fig. 2d, 2f). These findings suggest that liver- and adipose-derived sEVs may be involved in regulating metabolic improvements following VSG.

**Figure 3.**
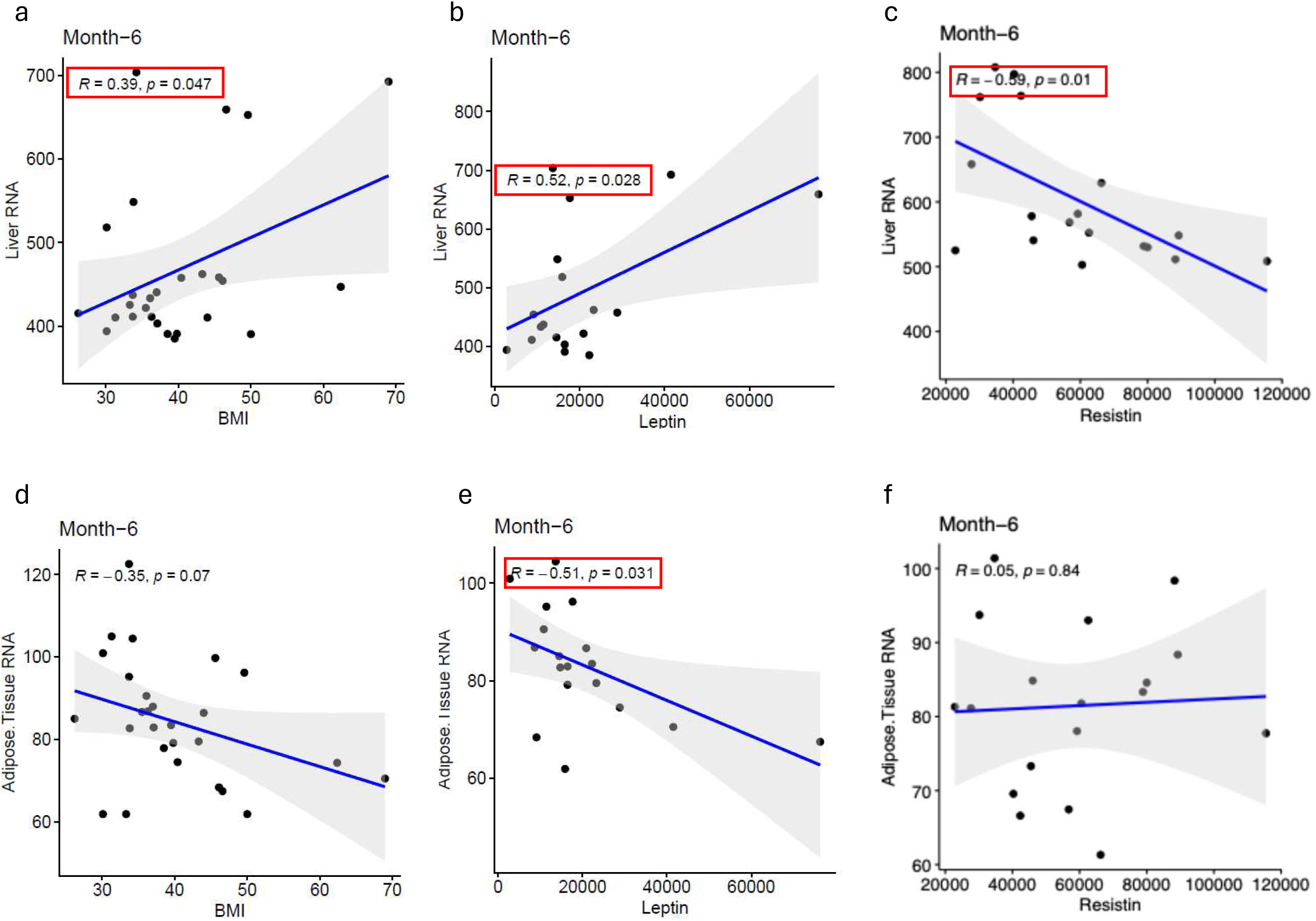
Correlation of RNA cargo of liver- and adipose tissue-specific genes within circulating sEVs with BMI, Leptin, and Resistin post-VSG. Counts of RNA cargo from liver-(a-c) and adipose tissue-(d-f) specific genes were analyzed for correlations with BMI (a, d), Leptin (b, e), and Resistin (c, f) at 6 months post-VSG.

### Differential Effects of Adipose and Liver sEVs on Metabolism and Liver Function Post-VSG

To further investigate the roles of liver- and adipose-derived sEVs, we performed a correlation analysis with HOMA-IR (Homeostatic Model Assessment of Insulin Resistance), a key indicator of insulin resistance that decreases following VSG (Fig. 4a). No significant correlations were observed between Liver- or Adipose-RNA cargo levels and HOMA-IR, either pre- or post-surgery (Supplemental Fig. 3). We then analyzed the delta (Δ) values, calculated as the difference between 6-month post-surgery and pre-surgery measurements, to assess how changes in RNA cargo levels correlate with improvements in insulin resistance. ΔHOMA-IR values were positively correlated with ΔAdipose-RNA cargo, suggesting that adipose-derived sEVs may contribute to improved insulin sensitivity following VSG. In contrast, no significant correlation was observed between ΔLiver-RNA cargo and ΔHOMA-IR (Fig. 4b, 4c). A similar pattern was observed with ALT, a marker of liver damage that decreased post-VSG (Fig. 4d). ΔALT was not correlated with ΔLiver-RNA cargo (Fig. 4e) but showed a positive correlation with ΔAdipose-RNA cargo (Fig. 4f). These results suggest a specific involvement of adipose-derived sEVs in enhancing both insulin sensitivity and liver function following VSG, while liver-derived sEVs may play a distinct, non-overlapping metabolic role.

**Figure 4.**
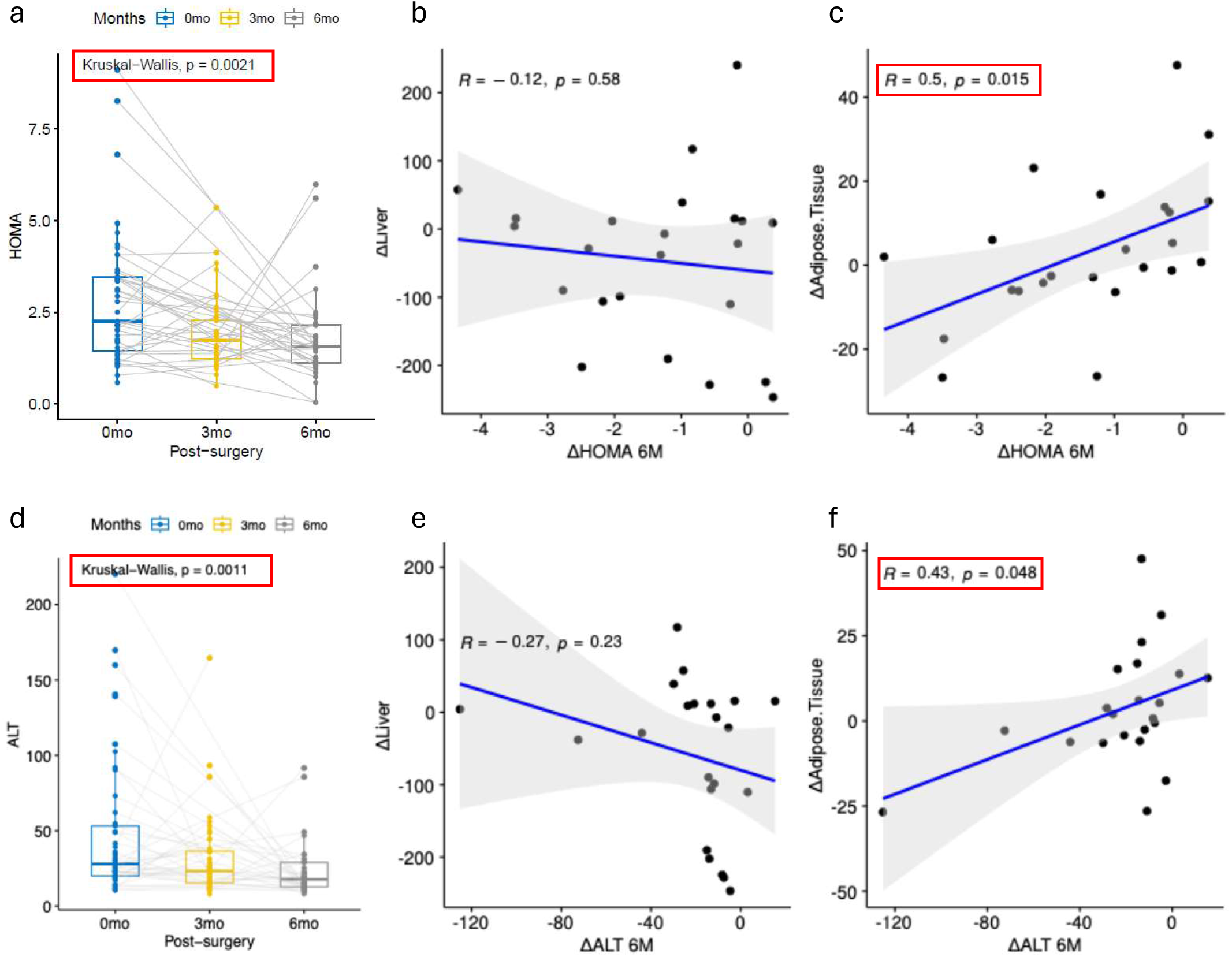
Changes in HOMA-IR and ALT and their correlation with Delta RNA cargo counts of liver- and adipose tissue-specific genes. (a) HOMA-IR in obese adolescents measured pre-surgery (n=58), at 3 months (n=44), and 6 months (n=40) post-VSG. (b, c) Correlation analyses of Delta HOMA-IR (6 months post-VSG minus pre-surgery) with Delta RNA cargo counts (6 months post-VSG minus pre-surgery) of liver-(b) and adipose tissue-(c) specific genes. (d) ALT levels in obese adolescents measured pre-surgery (n=58), at 3 months (n=44), and 6 months (n=40) post-VSG. (e, f) Correlation analyses of Delta ALT (6 months post-VSG minus pre-surgery) with Delta RNA cargo counts (6 months post-VSG minus pre-surgery) of liver-(e) and adipose tissue-(f) specific genes.

### Liver-Derived EV-RNA Links to BCAA Metabolism

Since adipose-derived sEVs appear to be associated with liver function, but liver-derived sEVs are not, we explored whether liver-derived sEVs might be linked to parameters related to other organs, particularly adipose tissue. This is relevant because Liver-RNA cargo was associated with leptin and resistin levels, both of which are predominantly secreted by adipose tissue. Branched-chain amino acids (BCAAs), which are associated with obesity and diabetes, showed a reduction post-VSG (Fig. 5a, Supplemental Fig. 4). Notably, changes in BCAA levels (ΔBCAA), particularly ΔValine and ΔLeucine, were positively correlated with ΔLiver-RNA cargo (Fig. 5b, Supplemental Fig. 4b and 4e), but no such correlation was found with ΔAdipose-RNA cargo (Fig. 5c, Supplemental Fig. 4c and 4f). These findings suggest that liver-derived sEVs may regulate systemic metabolism through associations with BCAA metabolism and adipose tissue-secreted hormones, highlighting distinct inter-organ communication pathways via sEVs post-VSG (Fig 7).

**Figure 5.**
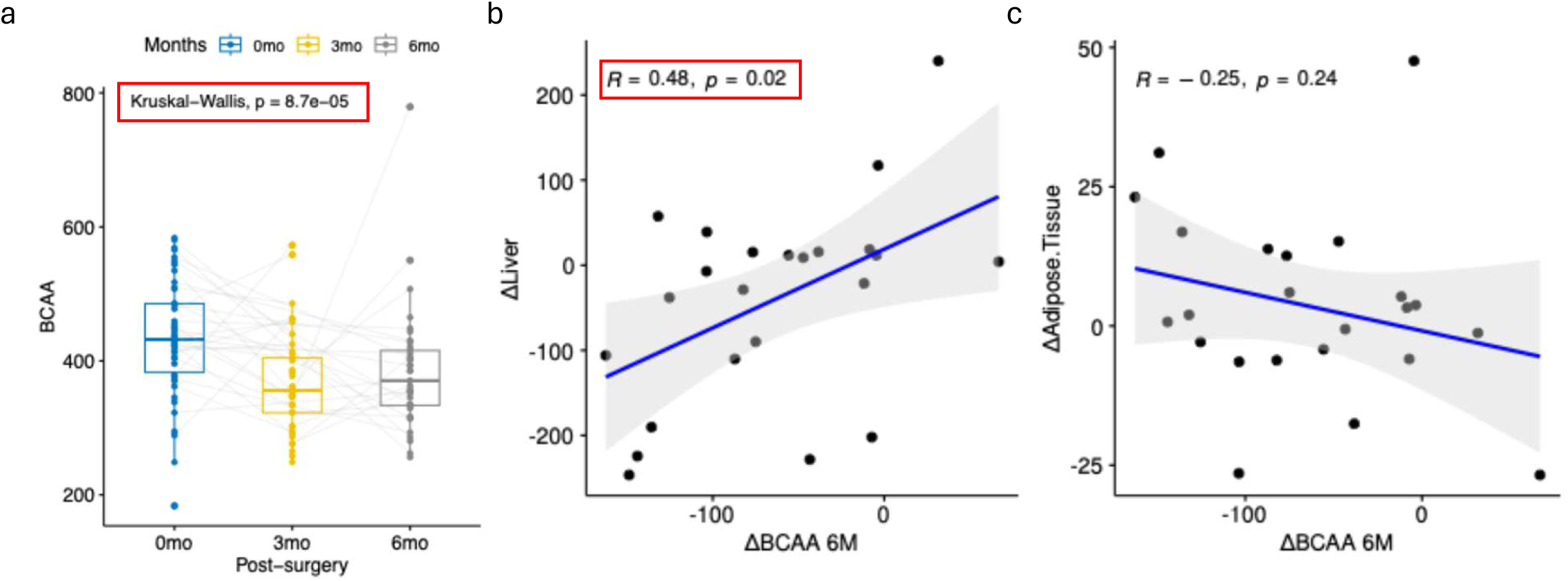
Changes in BCAA and their correlation with Delta RNA cargo counts of liver- and adipose tissue-specific genes. (a) Levels of circulating BCAA in obese adolescents measured pre-surgery (n=58), 3 months (n=44), and 6 months (n=40) post-VSG. (b, c) Correlation analyses of Delta BCAA (6 months post-VSG minus pre-surgery) with Delta RNA cargo counts (6 months post-VSG minus pre-surgery) of liver-(b) and adipose tissue-(c) specific genes.

## Discussion

Bariatric surgery promotes robust weight loss and rapid and dramatic metabolic improvements in patients with obesity [1–4], with VSG being the most frequently performed bariatric procedure in adolescents due to its effectiveness and favorable safety profile [4]. Growing evidence suggests that these metabolic benefits result from enhanced inter-organ communication, with several molecular mechanisms identified as contributors, including alterations in the bile acid profile, gut microbiome, hormones, and adipokines [5, 16, 41–47]. However, the role of sEVs, particularly their RNA cargo, remains underexplored despite their emerging importance as mediators of inter-organ communication and metabolic regulation. Research into sEVs is essential for understanding how VSG drives systemic metabolic changes. Small EVs transport bioactive molecules, such as RNA, facilitating crosstalk between distant tissues, thereby influencing metabolic functions. Investigating the RNA cargo within key metabolic tissue-derived sEVs, such as liver and adipose tissue, offers valuable insights into how VSG modulates organ-specific and systemic metabolism.

To our knowledge, this is the first report describing a reduction in Liver-RNA cargo in sEVs following VSG in adolescents with obesity. Although previous studies have reported a reduction in liver-derived sEVs following VSG [35, 36], our findings suggest that VSG influences sEV secretion and RNA cargo profiles in a tissue-specific manner, with distinct metabolic effects on the liver and adipose tissue. Notably, the sEV mRNA cargo representing other tissues, such as skeletal muscle and the gastrointestinal tract, did not show a similar trend in this cohort. These data indicate that the liver and adipose tissue are key metabolic organs responding to VSG by modulating its sEV cargo system in adolescents with obesity.

One of the key findings from our study was the emergence of statistically significant correlations between Liver-RNA cargo and Adipose-RNA cargo with key metabolic parameters, including BMI, leptin, and resistin, at 6 months post-VSG, which were absent at baseline. Notably, while VSG did not have a robust effect on Adipose-RNA cargo levels (Fig. 2f), a significant correlation between Adipose-RNA cargo and leptin levels was observed (Fig. 3e). This result suggests that severe obesity disrupts interactions between RNA cargo of sEV and metabolic parameters, and VSG may help restore these molecular interactions. The re-establishment of these interactions post-VSG may play a crucial role in the observed metabolic improvements. This highlights that VSG not only addresses metabolic dysfunction at the tissue level but also enhances inter-organ communication, which is essential for maintaining metabolic homeostasis.

Additionally, our observation that changes in Adipose-RNA cargo (ΔAdipo-RNA cargo) between pre- and post-VSG positively correlated with change in HOMA-IR and ALT underscores the importance of adipose-derived EVs in the post-VSG metabolic recovery. Although the overall levels of Adipose-RNA cargo in sEVs remained relatively unchanged after VSG (Fig. 2f), the significant correlations between ΔAdipo-RNA cargo and these metabolic parameters highlight the functional relevance of adipose-derived sEVs. This finding suggests that adipose-derived sEVs may contribute to rapid improvement of insulin resistance[8, 48, 49], a major risk factor for T2D, and enhanced liver function following VSG [4].

Individuals with obesity, insulin resistance, and T2D tend to have higher plasma BCAA levels compared to those who are metabolically healthy [50, 51]. Abnormal BCAA catabolism, resulting in BCAA accumulation, has been shown to induce insulin resistance [52–54]. A recent meta-analysis confirmed that bariatric surgery reduces plasma BCAA levels, contributing to metabolic improvements [55]. Skeletal muscle is the primary site of BCAA catabolism, with adipose tissue also playing a role in BCAA-related metabolic regulation. Impaired BCAA breakdown in both skeletal muscle and adipose tissue contributes to elevated circulating BCAA levels, which are associated with insulin resistance and metabolic disorders [52–54]. Emerging research further suggests that dysfunction in adipose tissue BCAA metabolism may exacerbate insulin resistance and other metabolic diseases [52]. Our findings suggest that liver-derived sEVs may influence BCAA metabolism in skeletal muscle and adipose tissue, or alternatively, BCAA levels may regulate sEV biogenesis in the liver. Further investigation will be needed to elucidate the interplay between liver-derived sEVs and BCAA metabolism and their combined role in systemic metabolic regulation.

In our study, the positive correlation between changes in Liver-RNA cargo (ΔLiver-RNA cargo) and changes in BCAA levels, with no similar association for ΔAdipo-RNA cargo, highlights the liver’s distinct role in BCAA metabolism post-VSG. These findings suggest that inter-organ communication via the sEV system plays a critical role in modulating insulin resistance and BCAA metabolism, contributing to the metabolic improvements observed after surgery (Fig 6). Together, these results emphasize the complementary roles of liver- and adipose-derived sEVs in post-VSG metabolic recovery. The liver appears to regulate BCAA metabolism, while adipose tissue contributes to improvements in insulin sensitivity. This coordinated response through the sEV system suggests that VSG restores molecular interactions and promotes metabolically healthy physiology, both at the tissue level and through inter-organ communication.

**Figure 6.**
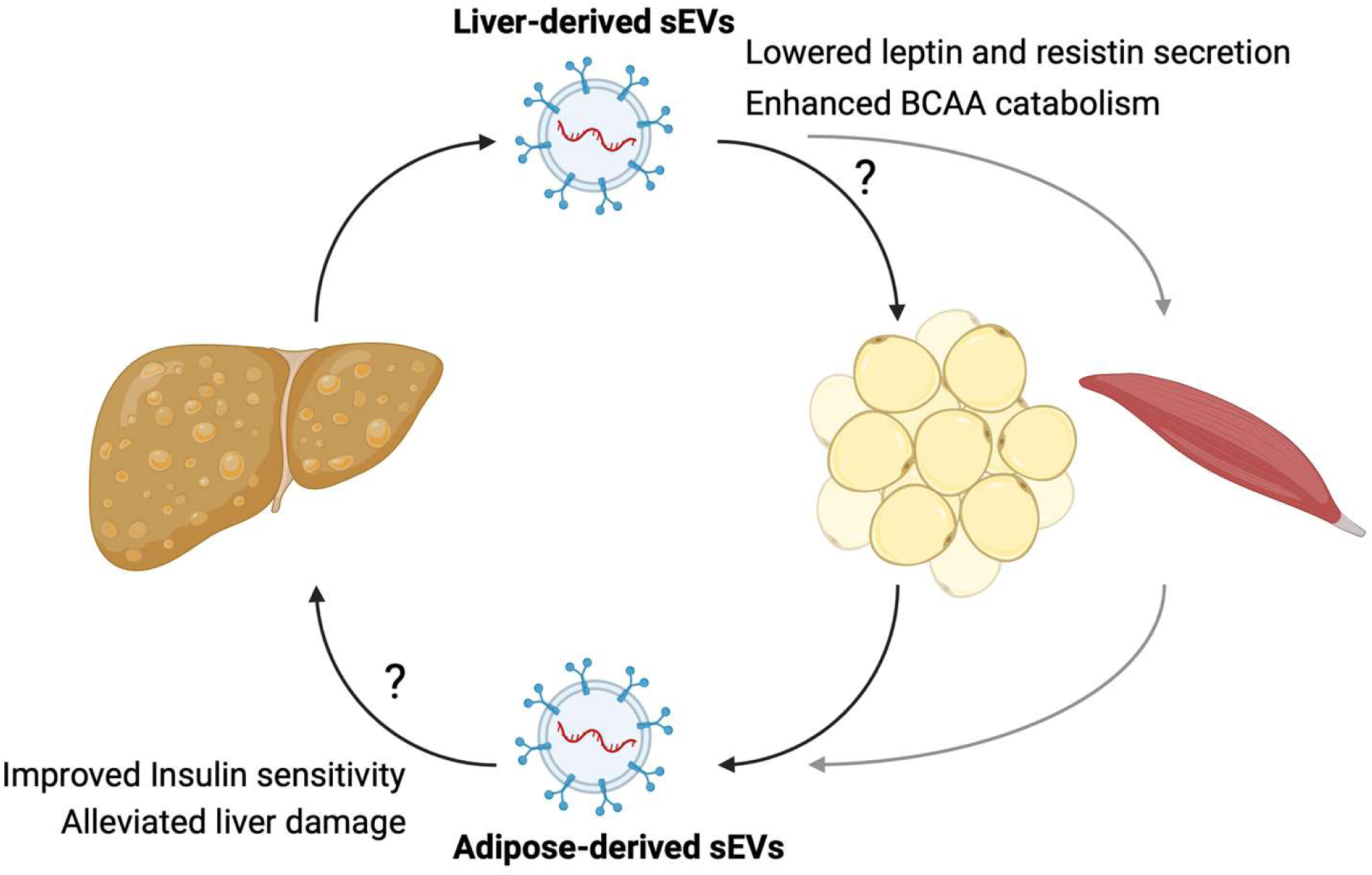
Proposed effects of bariatric surgery on liver- and adipose-derived sEVs and their metabolic outcomes. Bariatric surgery may modulate the release of small extracellular vesicles (sEVs) from adipose tissue and liver, influencing systemic metabolic outcomes. Liver-derived sEVs are proposed to improve insulin sensitivity, enhance BCAA catabolism, and regulate leptin and resistin secretion. Adipose tissue-derived sEVs may contribute to improved insulin sensitivity and alleviated liver damage. The exact mechanisms by which these sEVs drive post-surgery metabolic improvements remain to be fully elucidated (indicated by question marks).

The strengths of this study include a relatively large cohort of adolescents with obesity undergoing bariatric surgery, the analyses of tissue-specific RNAs in circulating sEVs, and the ability to capture longitudinal changes in metabolic parameters. However, several limitations should be noted. Inconsistent sample availability across time points (0 month n=58, 3 month n=45, 6 month n=41), and the absence of dietary history (pre-op versus post-op) may have affected the interpretation of changes in sEV profile and clinical parameters. Additionally, this study was conducted at a single center, and the observational nature of the data limits the ability to confirm causal relationships.

In conclusion, our study provides novel evidence that VSG modulates the sEV cargo of both liver and adipose tissue, and restore molecular interactions that are essential for metabolic improvements. These findings underscore the importance of sEV-mediated inter-organ communication in metabolic regulation and highlight the distinct roles of liver- and adipose-derived sEVs to BCAA metabolism and insulin resistance. Future research should investigate the functional significance of these interactions to identify new therapeutic strategies to correct metabolic dysfunction in obesity.

## Abbreviations

ALT: (Alanine aminotransferase)
BCAA: (branched chains amino acids)
BMI: (body mass index)
HOMA-IR: (Homeostatic Model Assessment of Insulin Resistance)
HPA: (Human Protein Atlas)
MASH: (metabolic-dysfunction associated steatotic hepatitis)
MASLD: (metabolic-dysfunction associated steatotic liver disease)
sEVs: (small extracellular vesicles)
T2D: (type 2 diabetes)
VSG: (vertical sleeve gastrectomy)

## Acknowledgments

We would like to acknowledge the clinical research coordinators for the Pediatric Diabetes and Obesity Center at Cincinnati Children’s Hospital Medical center for their support on this project.

## Data availability

Datasets are presented in the main manuscript, or Datasets are available upon request to the corresponding author.

## Authors’ relationships and activities

no relationships or activities that have influenced our work Contribution Statement: Each author listed had substantial contributions to conception and design, acquisition of data or analysis and interpretation of data, drafted the article or reviewed it critically for important intellectual content, and gave final approval of the version to be published.

---

**Supplemental Figure 1.**
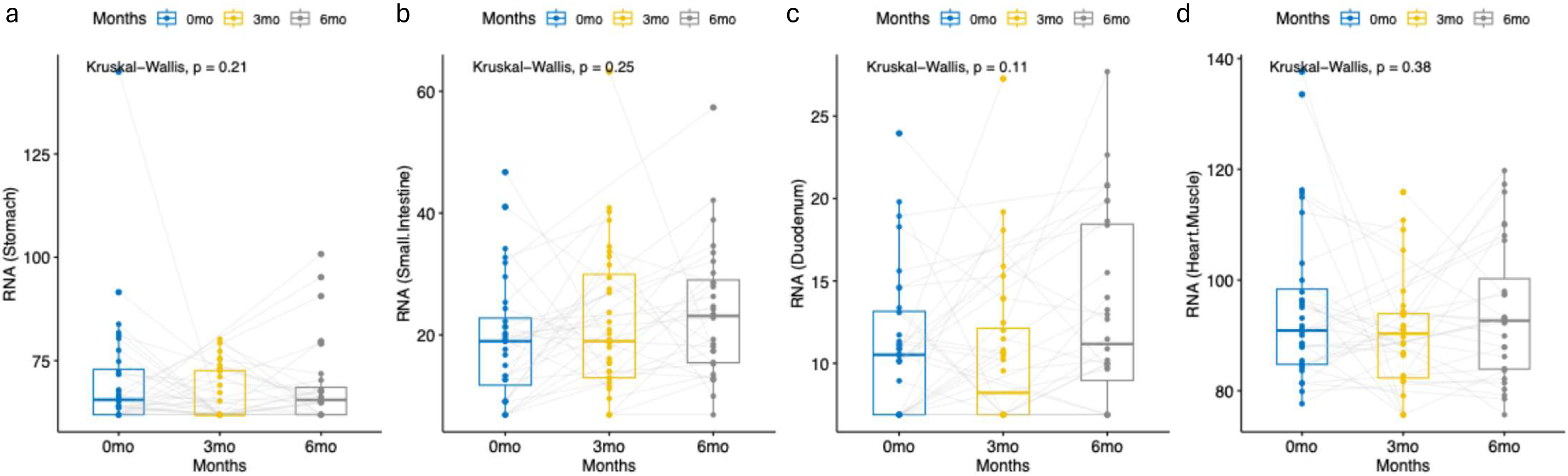
Tissue-specific genes’ cargo within circulating sEVs pre- and post-VSG. (a-d) RNA counts of tissue-specific genes within circulating sEVs for stomach (a), small intestine (b, duodenum (c) and heart (d).

**Supplemental Figure 2.**
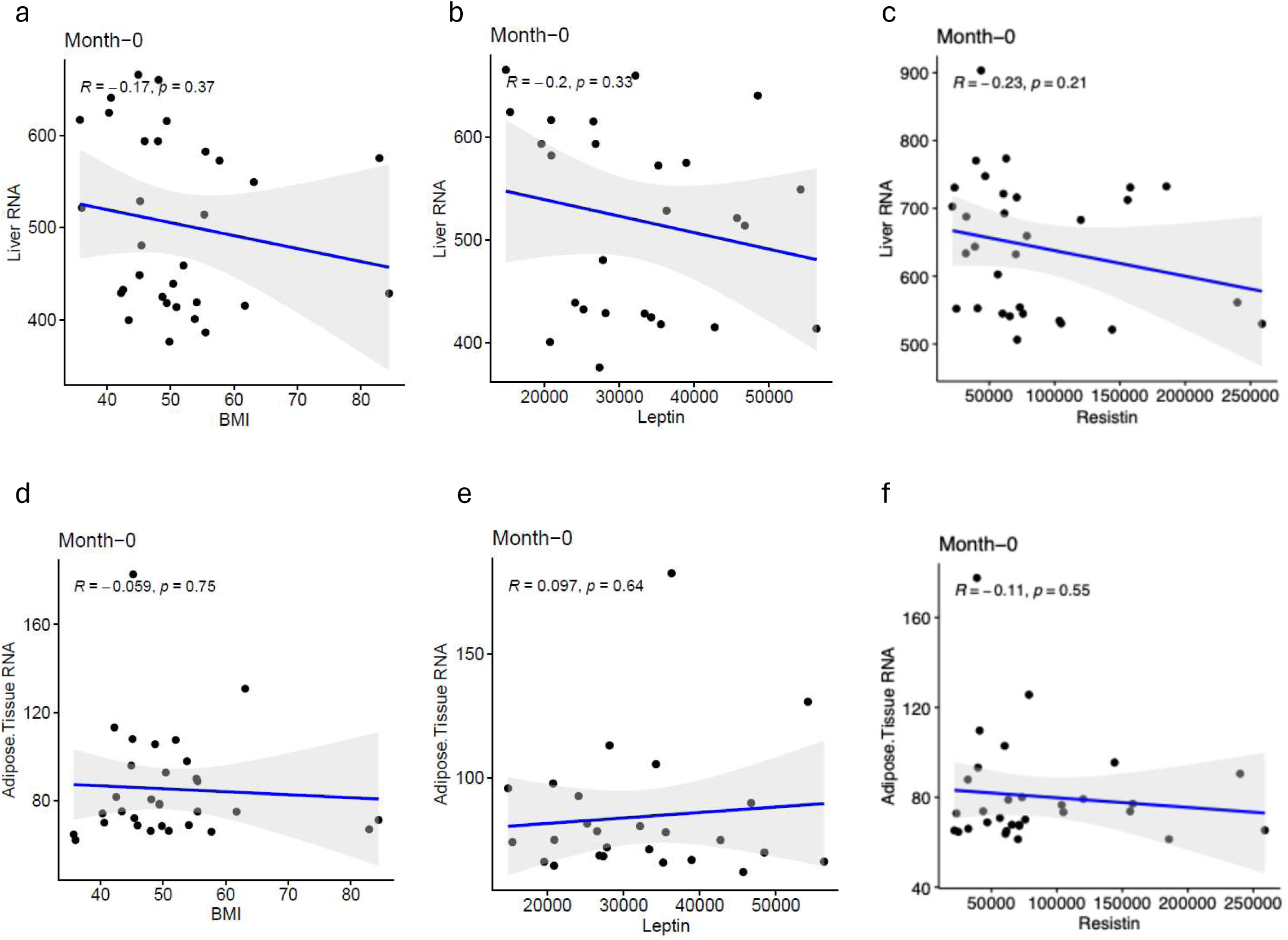
Correlation of RNA cargo of liver- and adipose tissue-specific genes with BMI, Leptin, and Resistin pre-VSG. Counts of RNA cargo from liver-(a-c) and adipose tissue-(d-f) specific genes were analyzed for correlations with BMI (a, d), Leptin (b, e), and Resistin (c, f) pre-VSG.

**Supplemental Figure 3.**
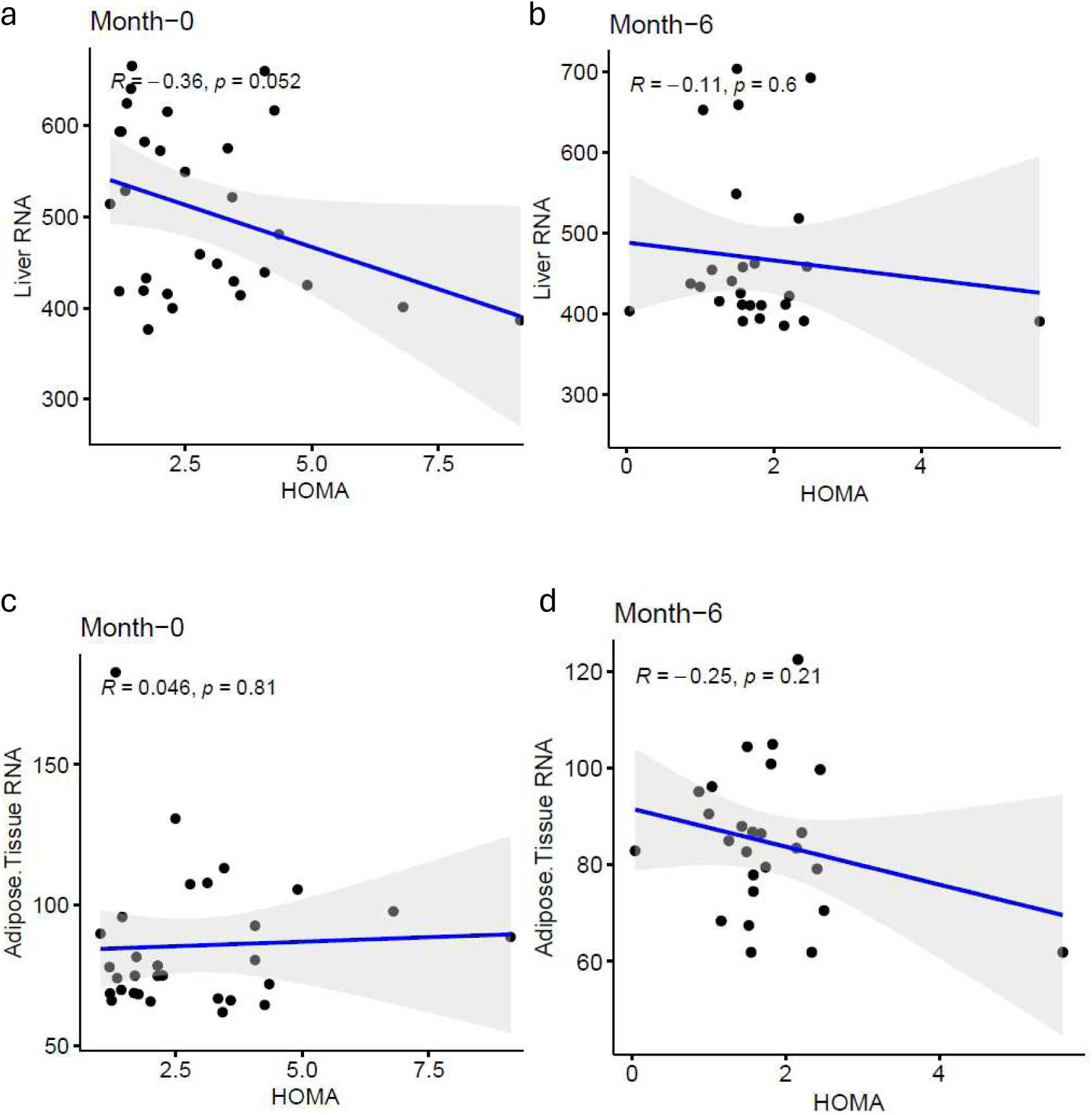
Correlation of RNA cargo of liver- and adipose tissue-specific genes with HOMA-IR pre- and post-VSG. Counts of RNA cargo from liver-(a, b) and adipose tissue-(c, d) specific genes were analyzed for correlations with HOMA-IR pre-(a, c) and post- (b, d) VSG.

**Supplementary Figure 4.**
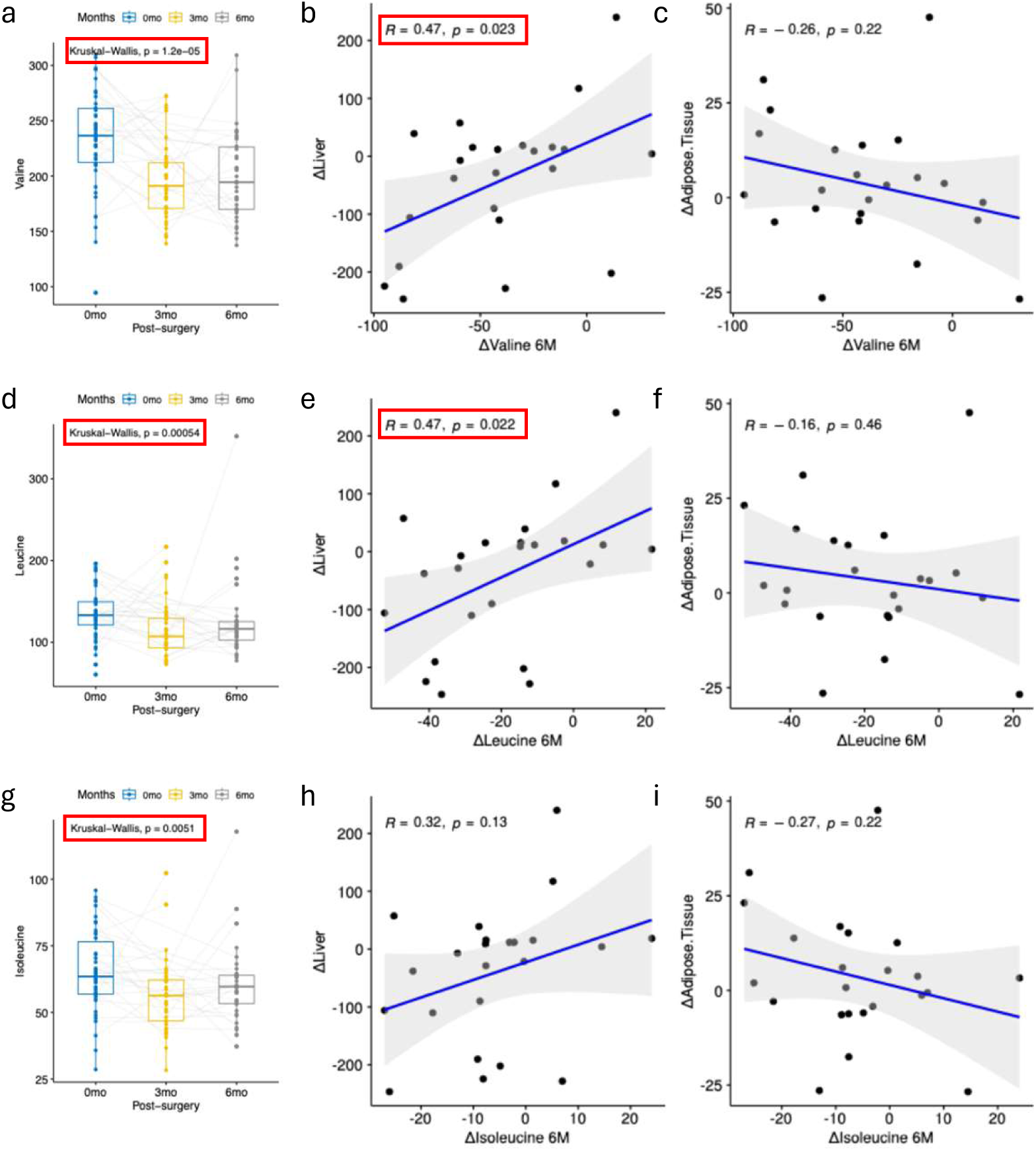
Changes in individual BCAAs and their correlation with Delta RNA cargo counts of liver- and adipose tissue-specific genes. (a, d, g) Levels of circulating valine (a), leucine (d), and isoleucine (g) in obese adolescents measured pre-surgery (n=58), 3 months (n=44), and 6 months (n=40) post-VSG. (b, c, e, f, h, i) Correlation analyses of Delta values (6 months post-VSG minus pre-surgery) for valine (b, c), leucine (e, f), and isoleucine (h, i) with Delta RNA cargo counts of liver-(b, e, h) and adipose tissue-(c, f, i) specific genes.

## References

1. Hampl, S.E., et al., Clinical Practice Guideline for the Evaluation and Treatment of Children and Adolescents With Obesity. Pediatrics, 2023. 151(2).

2. Pratt, J.S.A., et al., ASMBS pediatric metabolic and bariatric surgery guidelines, 2018. Surg Obes Relat Dis, 2018. 14(7): p. 882–901.

3. Eisenberg, D., et al., 2022 American Society of Metabolic and Bariatric Surgery (ASMBS) and International Federation for the Surgery of Obesity and Metabolic Disorders (IFSO) Indications for Metabolic and Bariatric Surgery. Obes Surg, 2023. 33(1): p. 3-14.

4. Inge, T.H., A.P. Courcoulas, and M.A. Helmrath, Five-Year Outcomes of Gastric Bypass in Adolescents as Compared with Adults. Reply. N Engl J Med, 2019. 381(9): p. e17.

5. Ji, Y., et al., Effect of Bariatric Surgery on Metabolic Diseases and Underlying Mechanisms. Biomolecules, 2021. 11(11).

6. Ryder, J.R., et al., Ten-Year Outcomes after Bariatric Surgery in Adolescents. N Engl J Med, 2024. 391(17): p. 1656–1658.

7. Swertfeger, D., et al., Presurgery health influences outcomes following vertical sleeve gastrectomy in adolescents. Obesity (Silver Spring), 2024. 32(6): p. 1187–1197.

8. Buchwald, H., et al., Bariatric surgery: a systematic review and meta-analysis. JAMA, 2004. 292(14): p. 1724–37.

9. Schauer, P.R., et al., Effect of laparoscopic Roux-en Y gastric bypass on type 2 diabetes mellitus. Ann Surg, 2003. 238(4): p. 467–84; discussion 84-5.

10. Ghanem, O.M., et al., Continued Diabetes Remission Despite Weight Recurrence: Gastric Bypass Long-Term Metabolic Benefit. J Am Coll Surg, 2024. 238(5): p. 862–871.

11. Courcoulas, A.P., et al., Long-Term Outcomes of Medical Management vs Bariatric Surgery in Type 2 Diabetes. JAMA, 2024. 331(8): p. 654–664.

12. Beckman, L.M., T.R. Beckman, and C.P. Earthman, Changes in gastrointestinal hormones and leptin after Roux-en-Y gastric bypass procedure: a review. J Am Diet Assoc, 2010. 110(4): p. 571–84.

13. Terra, X., et al., Long-term changes in leptin, chemerin and ghrelin levels following different bariatric surgery procedures: Roux-en-Y gastric bypass and sleeve gastrectomy. Obes Surg, 2013. 23(11): p. 1790–8.

14. Korner, J., et al., Effects of Roux-en-Y gastric bypass surgery on fasting and postprandial concentrations of plasma ghrelin, peptide YY, and insulin. J Clin Endocrinol Metab, 2005. 90(1): p. 359–65.

15. Geloneze, B., et al., Serum leptin levels after bariatric surgery across a range of glucose tolerance from normal to diabetes. Obes Surg, 2001. 11(6): p. 693–8.

16. Sebunova, N., et al., Changes in adipokine levels and metabolic profiles following bariatric surgery. BMC Endocr Disord, 2022. 22(1): p. 33.

17. Reifenberg, P. and A. Zimmer, Branched-chain amino acids: physico-chemical properties, industrial synthesis and role in signaling, metabolism and energy production. Amino Acids, 2024. 56(1): p. 51.

18. Nie, C., et al., Branched Chain Amino Acids: Beyond Nutrition Metabolism. Int J Mol Sci, 2018. 19(4).

19. Fine, K.S., J.T. Wilkins, and K.T. Sawicki, Circulating Branched Chain Amino Acids and Cardiometabolic Disease. J Am Heart Assoc, 2024. 13(7): p. e031617.

20. Choi, B.H., S. Hyun, and S.H. Koo, The role of BCAA metabolism in metabolic health and disease. Exp Mol Med, 2024. 56(7): p. 1552–1559.

21. Bozadjieva Kramer, N., et al., The Role of Elevated Branched-Chain Amino Acids in the Effects of Vertical Sleeve Gastrectomy to Reduce Weight and Improve Glucose Regulation. Cell Rep, 2020. 33(2): p. 108239.

22. Chaudhary, P.K., S. Kim, and S. Kim, Shedding Light on the Cell Biology of Platelet-Derived Extracellular Vesicles and Their Biomedical Applications. Life (Basel), 2023. 13(6).

23. Zaborowski, M.P., et al., Extracellular Vesicles: Composition, Biological Relevance, and Methods of Study. Bioscience, 2015. 65(8): p. 783–797.

24. Couch, Y., et al., A brief history of nearly EV-erything - The rise and rise of extracellular vesicles. J Extracell Vesicles, 2021. 10(14): p. e12144.

25. Freeman, D.W., et al., Altered Extracellular Vesicle Concentration, Cargo, and Function in Diabetes. Diabetes, 2018. 67(11): p. 2377–2388.

26. Eguchi, A., et al., Circulating adipocyte-derived extracellular vesicles are novel markers of metabolic stress. J Mol Med (Berl), 2016. 94(11): p. 1241–1253.

27. Agouni, A., et al., Endothelial dysfunction caused by circulating microparticles from patients with metabolic syndrome. Am J Pathol, 2008. 173(4): p. 1210–9.

28. Javeed, N., Shedding Perspective on Extracellular Vesicle Biology in Diabetes and Associated Metabolic Syndromes. Endocrinology, 2019. 160(2): p. 399–408.

29. Huang-Doran, I., C.Y. Zhang, and A. Vidal-Puig, Extracellular Vesicles: Novel Mediators of Cell Communication In Metabolic Disease. Trends Endocrinol Metab, 2017. 28(1): p. 3–18.

30. Povero, D., et al., Circulating extracellular vesicles with specific proteome and liver microRNAs are potential biomarkers for liver injury in experimental fatty liver disease. PLoS One, 2014. 9(12): p. e113651.

31. Fu, S., et al., Extracellular vesicles in cardiovascular diseases. Cell Death Discov, 2020. 6: p. 68.

32. Kim, A., A.S. Shah, and T. Nakamura, Extracellular Vesicles: A Potential Novel Regulator of Obesity and Its Associated Complications. Children (Basel), 2018. 5(11).

33. Myronovych, A., et al., Intestinal extracellular vesicles are altered by vertical sleeve gastrectomy. Am J Physiol Gastrointest Liver Physiol, 2021. 320(2): p. G153–G165.

34. Witczak, J.K., et al., Bariatric Surgery Is Accompanied by Changes in Extracellular Vesicle-Associated and Plasma Fatty Acid Binding Protein 4. Obes Surg, 2018. 28(3): p. 767–774.

35. Nakao, Y., et al., Circulating extracellular vesicles are a biomarker for NAFLD resolution and response to weight loss surgery. Nanomedicine, 2021. 36: p. 102430.

36. Rega-Kaun, G., et al., Changes of Circulating Extracellular Vesicles from the Liver after Roux-en-Y Bariatric Surgery. Int J Mol Sci, 2019. 20(9).

37. Langevin, S.M., et al., Comparability of the small RNA secretome across human biofluids concomitantly collected from healthy adults. PLoS One, 2020. 15(4): p. e0229976.

38. Walsh, K.B., et al., miR-181a Mediates Inflammatory Gene Expression After Intracerebral Hemorrhage: An Integrated Analysis of miRNA-seq and mRNA-seq in a Swine ICH Model. J Mol Neurosci, 2021. 71(9): p. 1802–1814.

39. Pereira, S., et al., Tissue-Specific Effects of Leptin on Glucose and Lipid Metabolism. Endocr Rev, 2021. 42(1): p. 1–28.

40. Li, Y., et al., Resistin, a Novel Host Defense Peptide of Innate Immunity. Front Immunol, 2021. 12: p. 699807.

41. Browning, M.G., et al., Changes in Bile Acid Metabolism, Transport, and Signaling as Central Drivers for Metabolic Improvements After Bariatric Surgery. Curr Obes Rep, 2019. 8(2): p. 175–184.

42. Cole, A.J., et al., The Influence of Bariatric Surgery on Serum Bile Acids in Humans and Potential Metabolic and Hormonal Implications: a Systematic Review. Curr Obes Rep, 2015. 4(4): p. 441–50.

43. Ilhan, Z.E., et al., Temporospatial shifts in the human gut microbiome and metabolome after gastric bypass surgery. NPJ Biofilms Microbiomes, 2020. 6(1): p. 12.

44. Wahlstrom, A., et al., Alterations in bile acid kinetics after bariatric surgery in patients with obesity with or without type 2 diabetes. EBioMedicine, 2024. 106: p. 105265.

45. Arakawa, R., et al., Prospective study of gut hormone and metabolic changes after laparoscopic sleeve gastrectomy and Roux-en-Y gastric bypass. PLoS One, 2020. 15(7): p. e0236133.

46. Holst, J.J., et al., Mechanisms in bariatric surgery: Gut hormones, diabetes resolution, and weight loss. Surg Obes Relat Dis, 2018. 14(5): p. 708–714.

47. Meek, C.L., et al., The effect of bariatric surgery on gastrointestinal and pancreatic peptide hormones. Peptides, 2016. 77: p. 28–37.

48. Bikman, B.T., et al., Mechanism for improved insulin sensitivity after gastric bypass surgery. J Clin Endocrinol Metab, 2008. 93(12): p. 4656–63.

49. Bradley, D., et al., Gastric bypass and banding equally improve insulin sensitivity and beta cell function. J Clin Invest, 2012. 122(12): p. 4667–74.

50. Caballero, B., N. Finer, and R.J. Wurtman, Plasma amino acids and insulin levels in obesity: response to carbohydrate intake and tryptophan supplements. Metabolism, 1988. 37(7): p. 672–6.

51. Felig, P., E. Marliss, and G.F. Cahill, Jr., Plasma amino acid levels and insulin secretion in obesity. N Engl J Med, 1969. 281(15): p. 811–6.

52. Shah, H., et al., BCAAs acutely drive glucose dysregulation and insulin resistance: role of AgRP neurons. Nutr Diabetes, 2024. 14(1): p. 40.

53. White, P.J., et al., Insulin action, type 2 diabetes, and branched-chain amino acids: A two-way street. Mol Metab, 2021. 52: p. 101261.

54. Newgard, C.B., et al., A branched-chain amino acid-related metabolic signature that differentiates obese and lean humans and contributes to insulin resistance. Cell Metab, 2009. 9(4): p. 311–26.

55. Barati-Boldaji, R., et al., Bariatric surgery reduces branched-chain amino acids’ levels: a systematic review and meta-analysis. Nutr Res, 2021. 87: p. 80–90.

