## Supplemental Table 1 for "Exploring Organ-Specific Extracellular Vesicles in Metabolic Improvements Following Bariatric Surgery in Adolescents with Obesity"

| Liver | Adipose tissu | Skeletal Mus | Stomach | Small intesti | Heart | Duodenum |
|---|---|---|---|---|---|---|
| ASGR1 | AC009477.2 | AC083902.2 | A4GNT | CPO | ACTC1 | ADA |
| ASGR2 | ACACB | ACTA1 | ANXA10 | FABP6 | ANKRD1 | MLN |
| A1BG | ACVR1C | ACTN3 | ATP4A | KCNJ13 | BMP10 | PDX1 |
| ABCB11 | ADIPOQ | ADSSL1 | ATP4B | NA | CASQ2 | S100G |
| ABCB4 | AL845331.2 | ALDOA | BARX1 | NAALADL1 | CCDC141 | TMPRSS15 |
| ACOT12 | AQP7 | AMPD1 | CAPN8 | P2RY4 | CHRNE | |
| ACSM5 | CFD | ANK1 | DPCR1 | | FABP3 | |
| ADH1A | CIDEA | ANKRD23 | GAGE12G | | LRRC10 | |
| ADH4 | CIDEC | ART1 | GAST | | MT1HL1 | |
| ADH6 | FABP4 | ASB10 | GHRL | | MYBPC3 | |
| AFM | GYG2 | ASB12 | GIF | | MYBPHL | |
| AGXT | KIF25 | ASB16 | GKN1 | | MYH6 | |
| AHSG | LEP | ATP1B4 | GKN2 | | MYL4 | |
| AKR1C4 | LGALS12 | ATP2A1 | KCNE2 | | MYL7 | |
| AKR1D1 | LIPE | CA3 | LIPF | | MYOZ2 | |
| ALB | PLIN1 | CACNA1S | MUC5AC | | NPPA | |
| AMBP | PLIN4 | CACNG1 | PGA3 | | NPPB | |
| ANG | PNPLA2 | CALML6 | PGA4 | | PLN | |
| ANGPTL3 | TIMP4 | CASQ1 | PGA5 | | PPP1R1C | |
| APCS | TUSC5 | CDH15 | PGC | | RD3L | |
| APOA2 | | CHRNA10 | RFLNA | | RYR2 | |
| APOA5 | | CHRND | SLC9A4 | | SBK2 | |
| APOC1 | | CHRNG | TFF1 | | SBK3 | |
| APOC2 | | CKM | | | SCN5A | |
| APOC3 | | CLCN1 | | | TECRL | |
| APOC4 | | COQ8A | | | TNNI3 | |
| APOC4-APOC2 | | DUPD1 | | | TNNI3K | |
| APOF | | DWORF | | | TNNT2 | |
| APOH | | ENO3 | | | | |
| ARID3C | | FBP2 | | | | |
| ASGR1 | | FEM1A | | | | |
| ASGR2 | | FGF6 | | | | |
| BAAT | | FGF8 | | | | |
| BDH1 | | FHL3 | | | | |
| BX248415.1 | | IDI2 | | | | |
| C5 | | JPH1 | | | | |
| C8A | | JSRP1 | | | | |
| C8B | | KBTBD13 | | | | |
| C9 | | KCNA7 | | | | |
| CCL16 | | KCNJ12 | | | | |
| CFB | | KIAA1024L | | | | |
| CFHR1 | | KLHL33 | | | | |

| | | |
|---|---|---|
| CFHR2 | | KLHL34 |
| CFHR3 | | KLHL40 |
| CFHR4 | | KLHL41 |
| CFHR5 | | LBX1 |
| CLEC1B | | LINC00116 |
| CP | | LRRC30 |
| CPB2 | | LSMEM1 |
| CPN1 | | MACROD1 |
| CPN2 | | MAPK12 |
| CYP1A2 | | MSS51 |
| CYP26A1 | | MUSTN1 |
| CYP2A6 | | MYADML2 |
| CYP2A7 | | MYBPC1 |
| CYP2B6 | | MYBPC2 |
| CYP2C8 | | MYF6 |
| CYP2C9 | | MYH1 |
| CYP2E1 | | MYH2 |
| CYP3A43 | | MYH4 |
| CYP3A7 | | MYL1 |
| CYP7A1 | | MYLK2 |
| CYP8B1 | | MYLPF |
| F12 | | MYOD1 |
| F13B | | MYOG |
| F2 | | MYOT |
| F7 | | MYOZ1 |
| F9 | | MYOZ3 |
| FCN2 | | NEB |
| FETUB | | OBSCN |
| FGA | | PHKG1 |
| FGB | | PPDPFL |
| FGF21 | | PPM1N |
| FGG | | PPP1R27 |
| FGL1 | | PRKAG3 |
| FMO3 | | PYGM |
| GBP7 | | RAPSN |
| GC | | RGS9BP |
| GCKR | | RPL3L |
| GDF2 | | RTN2 |
| GLS2 | | RYR1 |
| GYS2 | | SLN |
| HAMP | | SMTNL1 |
| HAO1 | | STAC3 |
| HGFAC | | SYPL2 |
| HP | | TMEM38A |
| HPR | | TMOD4 |

| | | |
|---|---|---|
| HPX | | TNNC2 |
| HRG | | TNNI1 |
| HSD17B13 | | TNNI2 |
| HSD17B6 | | TNNT1 |
| IL27 | | TNNT3 |
| INHBC | | TPM2 |
| INHBE | | TPM3 |
| ITIH1 | | TRIM72 |
| ITIH2 | | TTN |
| ITIH3 | | UCP3 |
| ITIH4 | | VGLL2 |
| KLKB1 | | YIPF7 |
| LEAP2 | | |
| LECT2 | | |
| LIPC | | |
| LPA | | |
| LRG1 | | |
| MASP2 | | |
| MAT1A | | |
| MBL2 | | |
| NR1I3 | | |
| OIT3 | | |
| ORM1 | | |
| ORM2 | | |
| PGLYRP2 | | |
| PLG | | |
| PLGLB1 | | |
| PLGLB2 | | |
| PON1 | | |
| PON3 | | |
| PRAMEF10 | | |
| PRG4 | | |
| PROC | | |
| PZP | | |
| RBP4 | | |
| RDH16 | | |
| RTP3 | | |
| SAA4 | | |
| SDS | | |
| SERPINA1 | | |
| SERPINA10 | | |
| SERPINA11 | | |
| SERPINA6 | | |
| SERPINA7 | | |
| SERPINC1 | | |

| |
|---|
| SERPIND1 |
| SERPINF2 |
| SLC10A1 |
| SLC13A5 |
| SLC17A2 |
| SLC22A1 |
| SLC22A10 |
| SLC22A25 |
| SLC22A9 |
| SLC25A47 |
| SLC27A5 |
| SLC2A2 |
| SLC38A3 |
| SLC38A4 |
| SLCO1B1 |
| SLCO1B3 |
| SPP2 |
| TAT |
| TDO2 |
| TF |
| TFR2 |
| TTC36 |
| TTPA |
| TTR |
| UGT1A3 |
| UGT1A4 |
| UGT2B10 |
| UGT2B4 |
| UROC1 |
| VTN |
